# A graph-based framework for multi-scale modeling of physiological transport

**DOI:** 10.1101/2021.09.14.460337

**Authors:** M. Deepa Maheshvare, Soumyendu Raha, Debnath Pal

**Affiliations:** Department of Computational and Data Sciences, Indian Institute of Science, Bangalore 560012, India

**Keywords:** biophysical interactions, discrete network model, functional networks, hierarchical modeling, microvasculature and microenvironment, solute transport, spatio-temporal dynamics, systems biology

## Abstract

Trillions of chemical reactions occur in the human body every second, where the generated products are not only consumed locally but also transported to various locations in a systematic manner to sustain homeostasis. Current solutions to model these biological phenomena are restricted in computability and scalability due to the use of continuum approaches where it is practically impossible to encapsulate the complexity of the physiological processes occurring at diverse scales. Here we present a discrete modeling framework defined on an interacting graph that offers the flexibility to model multiscale systems by translating the physical space into a metamodel. We discretize the graph-based metamodel into functional units composed of well-mixed volumes with vascular and cellular subdomains; the operators defined over these volumes define the transport dynamics. We predict glucose drift governed by advective-dispersive transport in the vascular subdomains of an islet vasculature and cross-validate the flow and concentration fields with finite-element based COMSOL simulations. Vascular and cellular subdomains are coupled to model the nutrient exchange occurring in response to the gradient arising out of reaction and perfusion dynamics. The application of our framework for modeling biologically relevant test systems shows how our approach can assimilate both multi-omics data from *in vitro* - *in vivo* studies and vascular topology from imaging studies for examining the structure-function relationship of complex vasculatures. The framework can advance simulation of whole-body networks at user-defined levels and is expected to find major use in personalized medicine and drug discovery.

**GRAPHICAL ABSTRACT:** 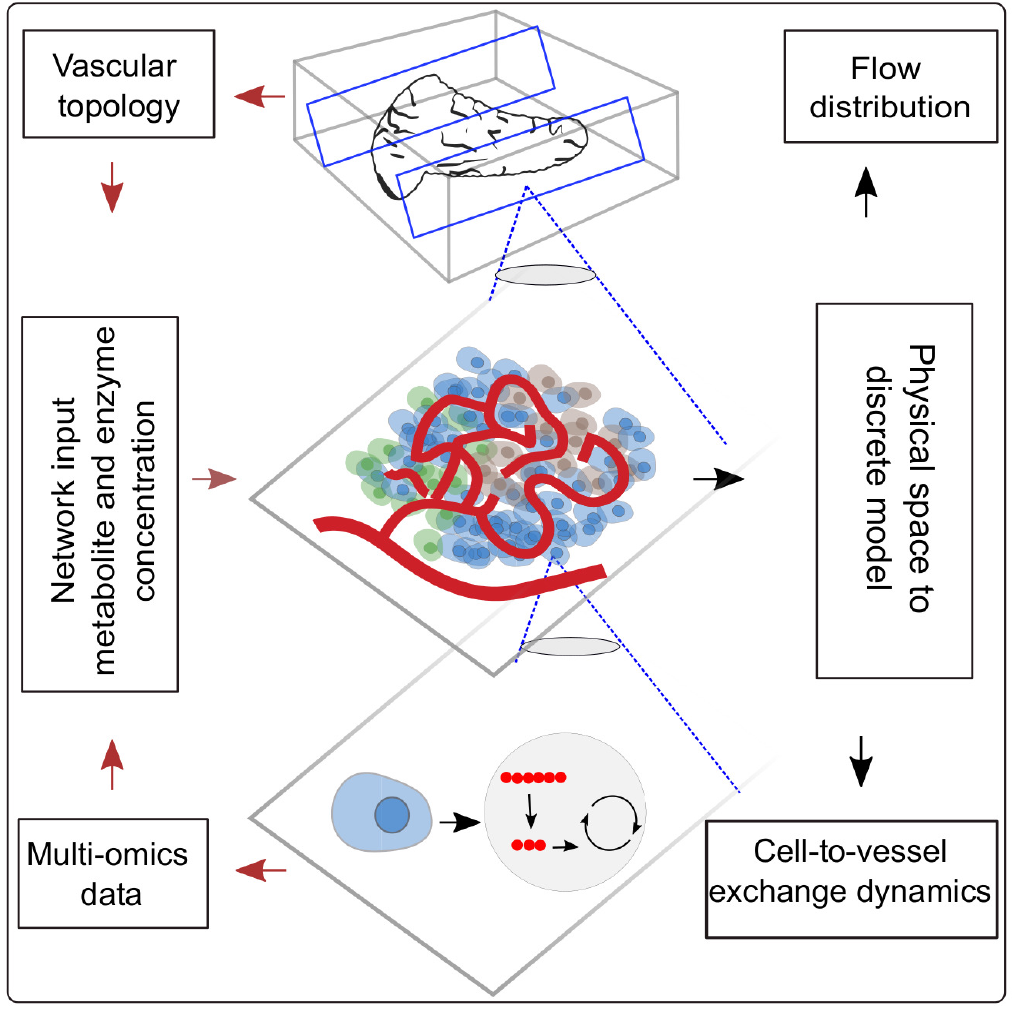

## 1 INTRODUCTION

The physiological system is a complex network in which each organ forms a subsystem and different subsystems interact to maintain overall homeostasis of the body. Within each subsystem functional networks exists at different levels of complexity. Metabolic and signaling networks within a cell, cell-to-cell communication networks in the extravascular region of tissue, cell-to-vessel communication networks, and the vascular network which couples the local dynamics to the global dynamics determine the functional behavior of all tissues. Bottom-up and top-down modeling approaches emulate the cellular dynamics and organ-level physiology. The ability to simultaneously capture the local and global dynamics by hierarchically bridging the communication networks existing across diverse scales is the key challenge in the holistic modeling of physiology.

Microscale cellular models use a bottom-up approach in which multi-omics data assimilated from high-throughput sources are employed to formulate and validate parameter-intensive kinetic models. These models capture the dependency of intracellular dynamics on metabolic steady states and flux distributions (König et al., 2012, Berndt et al., 2018a, Masid et al., 2020). Owing to cellular heterogeneity and the existence of intercellular communication, autocrine and paracrine signaling (Rao and Rizzo, 2020, Watts et al., 2016, Koh et al., 2012), the response elicited by a single cell cannot be scaled to a cell community. Cell population models, therefore, employ discrete modeling approaches for examining cell-to-cell interactions such as intra- and inter-islet synchronization established by gap junctional coupling (Pedersen et al., 2005, Barua and Goel, 2016).

Macroscopic organ scale compartment models (Sorensen, 1985) employ top-down modeling approach for predicting the bulk flow and elimination kinetics of biomolecules. These organ scale models rely on single-tube, parallel-tube, or tank-in-series approximations for idealizing distribution volume of blood into compartments (Gray and Tam, 1987). For improving the mechanistic understanding of tissue-vessel interaction, multiphase porous media-based models representing the tissue volume as intravascular and multi-region extravascular compartments (e.g. capillary-interstitial-parenchymal exchange unit) (Deussen and Bassingthwaighte, 1996, Chalhoub et al., 2007) emerged. However, these frameworks do not offer the possibility to fuse macro- and micro-scale models. Consequently, the effect of network architecture on microperfusion patterns (Dolenšek et al., 2015) and its influence on the nutrient exchange cannot be investigated by these compartment models.

To overcome the limitations of compartment-based models, continuum approaches have been put forth for understanding the implications of morphological changes on the functional response of an organ. In the extravascular domain, continuum approaches are helpful in estimating the collective response of a tissue mass where the bulk of the tissue is smeared and treated as a homogeneous domain. Although homogenization simplifies the complexity of the computational domain, the approach is limited in its ability to probe aspects such as the influence of heterogeneous arrangement of cells on nutrient release patterns. In the intravascular domain, continuum approaches are suitable for analyzing the effects of dilation of blood vessels, deformations that occur as a result of fluid-structure interactions. With the advancements in imaging studies, the availability of microvascular datasets offers the possibility to model large scale networks. However, discretizing the tortuous microvasculature vasculature for 3D modeling of the advection-dispersion physics gives rise to extensive computational overhead while employing continuum approaches. This has led to the development of graph- and hybrid-based approaches in which the vasculature infiltrating the tissue volume is represented as one dimensional network of pipes for modeling the flow and delivery of resources over networks (Beard and Bassingthwaighte, 2000, Fang et al., 2008, Heaton et al., 2012, Kojic et al., 2017, Safaei et al., 2018, Erlich et al., 2019). In summary, while efforts have been made to resolve: (i) spatial heterogeneity at subcellular scale (Ii et al., 2019), (ii) short-range communication in the microenvironment of cell communities, (iii) metabolic zonation in single sinusoid models (Berndt et al., 2018b), explicit models of long-range communications mediated by the vascular system remain underdeveloped at both intra-organ and inter-organ scale.

Towards this end we need a scalable hierarchical framework that allows us to bridge diverse scales for modeling production, consumption and distribution of biochemicals in a tissue microenvironment. We introduce a discrete modeling framework for simulating gradient-driven advection-dispersion-reaction physics of multispecies transport. Graph-theoretic approaches that have been proven successful in examining flow of information through large scale real-world networks is applied (Bellocchi and Geroliminis, 2020, Kumar et al., 2019, Besse and Faye, 2021) in this study. We resort to discrete-vector calculus and use the operators defined on a finite-graph to spatially discretize and formulate the transport dynamics in the vascular domain as a “tank-in-series” model. Further, the computational domain for establishing the vessel-to-cell exchange and cellular dynamics within the cell are set up by combining ideas from other multiscale and Krogh cylinder models (Berndt et al., 2018b, Frost et al., 2019). Dynamics of nutrient exchange from blood vessel to the layer of cells that lie in close proximity to the vessel surface is modeled; cellular reactions are explicitly modeled by representing cells as discrete volume nodes. Differential equations defining the interactions over nodal volumes embedded in the graph are solved by translating the physical domain into a metamodel in which the biophysical attributes are subsumed. This framework is suitable for the following key applications: (i) reduce the computational cost involved in the spatial discretization of large tissue volumes (Sec. 3.1.2); our discrete approach is geared towards obtaining fast solution by reducing the system dimension and the metamodel is scalable into any domain; (ii) probe the effect of flow topology on scalar transport and the sensitivity of concentration dynamics to network parameters and variations in physiological setpoints. (Sec. 3.2, 3.4); (iii) assimilate multi-omics data from *in vitro* and *in vivo* studies and vascular topology from imaging studies (Sec. 3.3) for examining the influence of structural changes on the functional response of a tissue. Our graph-based discrete modeling framework differs from the existing approaches in the following aspects. Conventional finite-difference, finite volume or finite-element based formulations operate on a continuous domain and the equations are discretized and approximate solutions are obtained. In our approach, we discretize the physical space and solve the equations on the graph which forms the discrete domain. This makes it possible to scale our framework to large networks; offers the flexibility to fuse multiscale models by encoding imaging data of vascular topology and omics data of cellular reactions to enhance systems level understanding.

The outline of this study is as follows. The procedure followed in translating the capillary vasculature into a weighted graph and the preliminary assumptions considered for setting up the computational domain are discussed in Section 2.1 and 2.2. The governing equations of the flow distribution and the mathematical formulation of the discrete model of advection-diffusion-reaction physics are presented (Sec. 2.3, 2.4, 2.5). We use our framework to model two physiologically relevant test systems: (i) advection-dispersion dynamics of glucose transport in the microvasculature, (ii) advection-dispersion-reaction dynamics of glucose-lactate exchange in the functionally coupled tissue-vascular domains (glucose-lactate dynamics is relevant in tumor metabolism where metabolic activity alters in tumor microenvironments (Yang et al., 2021) and in modeling fuel-stimulated insulin secretion (Jiang et al., 2007, Prentki et al., 2013)). By applying our method, we predict glucose drift in the islet vasculature and cross-validate the flow and concentration fields of the multiphysics simulation with COMSOL simulations (Sec. 3.1.1). We establish the cell-vessel link and predict the spatio-temporal evolution of glucose-lactate exchange in the extravascular and intravascular domains (Sec. 3.3). We test the model behavior for various flow topologies (Sec. 3.2), different pressure drops and glucose doses (Sec. 3.4). The network configurations illustrated in the applications presented in this work are the capillary blood vessels.

## 2 METHODOLOGY

For setting up the discrete-modeling framework to study the multiphysics coupling in multiscale systems, we start by introducing the steps involved in constructing the computational domain which is a metamodel of the physical space, dissection of the metamodel into subdomains which form the functional units and formulation of the mathematical operators. The three main steps involved in our workflow are illustrated in Figure 1. *Create skeleton*: The topological organization of the capillary network and biophysical characteristics such as length and diameter of the vessels in the network constitute the structural and anatomical characteristics relevant for setting up the computational domain. These characteristics are extracted in this step by skeletonizing the reconstructed vasculature (Sec. 2.1). *Solve flow distribution in the network*: The physical space is translated into a weighted graph representing a hydraulic circuit. The pressure and flow fields are computed over the network by establishing the relationship between the node and edge entities of the graph using Hagen-Poiseuille equation (Sec. 2.3). *Solve advection-dispersion-reaction dynamics*: The metamodel is subdivided into functional units composed of cell and vessel subdomains. We combine multiple scales by coupling the uptake and release flux of cell domain *ω*^*t*^ with the carrier-mediated exchange flux occurring at cell-vessel interface. The metabolic reactions occurring at the cellular scale are modeled by biochemical rate laws. And the mass transport in the capillary domain Ω^*bv*^ is described by coupling the cell-to-vessel influx or outflux with the gradient-driven advection-dispersion transport in the capillary domain Ω^*bv*^ (Sec. 2.4, 2.5). Henceforth, the superscripts *bv* and *t* denote the parameters and variables defined in the capillary blood vessel and tissue domains, respectively.

**Figure 1.**
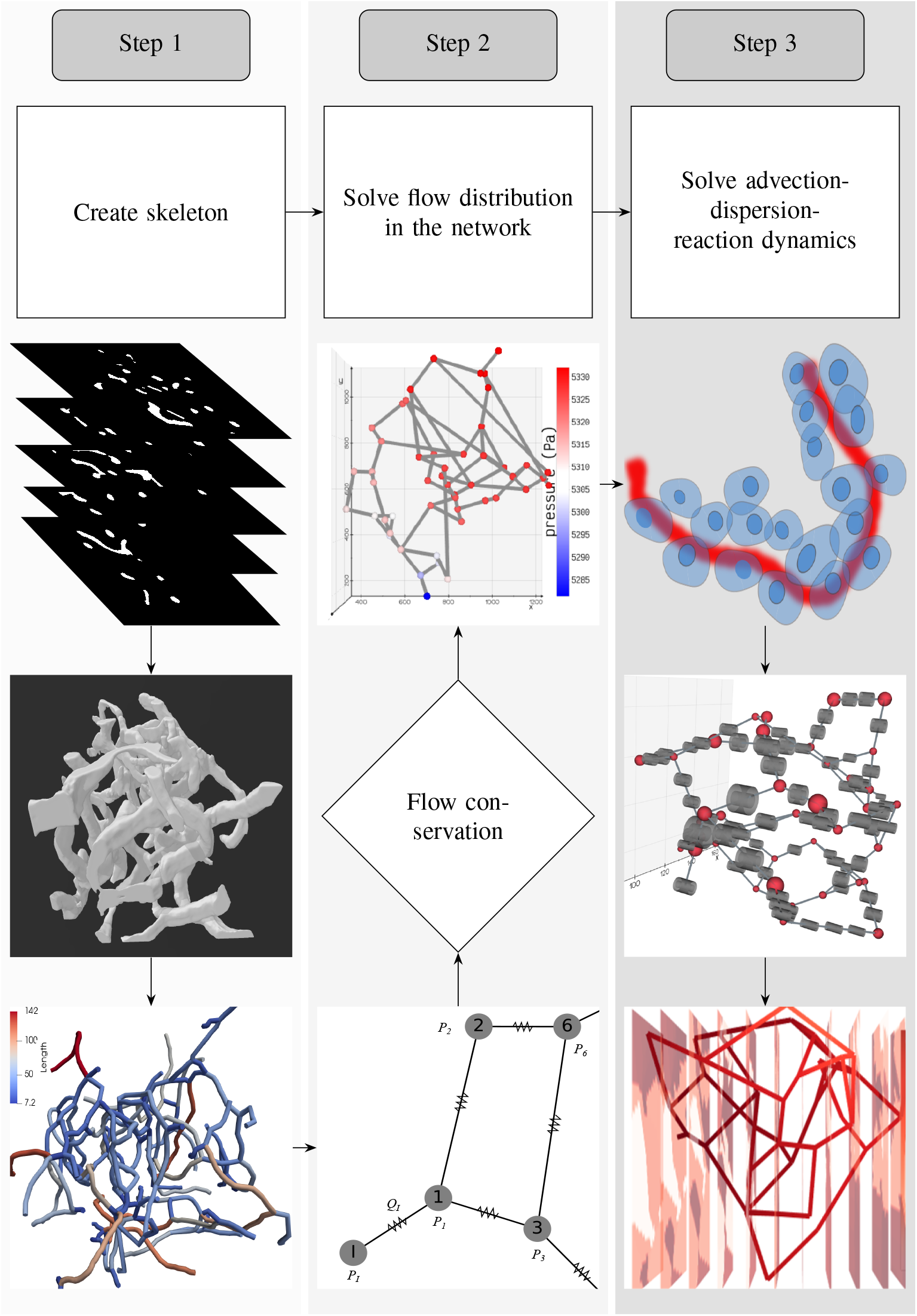
The workflow involved in setting up the system for simulating the spatio-temporal evolution of biochemical species is illustrated: *1. Create skeleton*: top, z-stack of microscopic images; middle, three dimensional volume of the reconstructed vascular geometry; bottom, the length of blood vessel branches are color-coded. *2. Solve flow distribution in the network* : bottom, the vascular network represented as a hydraulic circuit; middle: from the estimated nodal pressures, flow thorough the vessel branches are computed and checked whether the mass conservation holds at the junctions; top: 3D visualization of pressure gradient observed across the vasculature. *3. Solve advection-dispersion-reaction dynamics*: top: creation of the computational domain which includes blood vessel and the layer of the cells located in the vicinity of the outer surface of the capillary; middle: functional unit of the computational domain consisting of the vascular and cell sub-domains; bottom: concentration gradient observed across the vasculature after solving the advection-dispersion-reaction dynamics.

### 2.1 Construction of Capillary Networks

The topology of the blood vessels that exchange nutrients with the surrounding tissue is represented by one dimensional network of pipes. The biophysical attributes such as diameter and length of each segment, the blood vessel fragment connecting two branching points, are quantified as edge weights (𝕎) of the graph network (𝒢). We investigate four network configurations in this work. The 3D vasculature of pancreatic islet displayed in Figure 2**A** and both the configurations of tumor network shown in Figure 2**B**,**C** were reconstructed from binary images of vessels which were examined in Chen et al. (Chen et al., 2020). The tiff stack containing binary images were generated in their study by imaging the blood vessels, labeled with a fluorescent dye, using light sheet microscopy followed by segmentation of the vessels in *ilastik* (Sommer et al., 2011) toolkit which leverages a machine learning-based classification algorithm. We follow the workflow detailed below for generating a weighted graph from the binary images: (i) The multipage .tiff image was rendered into a 3D volume using the ray casting technique available in *3D Slicer* (Fedorov et al., 2012) by specifying the spacing of the image stack in the xyz directions. The dimensions of the input stack and domain size of the reconstructed volume are given in Table 1. (ii) The largest connected region of the segmented volume was filtered using the *Island* effect available in *Slicer* and the 3D object was exported in a stereolithography file (.stl) for skeletonization in the *Vascular Modelling Toolkit library (VMTK)* (Antiga et al., 2008). (iii) *vmtksurfaceclipper* was employed to open the surface at the network inlet and *vmtknetworkextraction* algorithm was utilized to skeletonize the geometry. This yielded a network with nodes (vertices) (𝕍) positioned at the N-furcation points or terminal ends, and edges (𝔼) formed by the vessel segment linking two nodes. (iv) Segment length *l*^*bv*^ and diameter *d*^*bv*^ of the blood vessels were computed by tracing the shortest path between two nodes and extracting the maximum inscribed sphere radius (thickness), respectively. The 2D structure of the mesentery vasculature displayed in Figure 2**D** was generated by parsing the diameter, length of vessels, and topology information available in *Amira* mesh file provided in Esposito et al (d’Esposito et al., 2018). Statistics of diameter and length distributions are shown in Figure 2**E**,**F**. We represent the skeletonized geometry of the capillaries as a weighted graph, 𝒢 (𝕍∈ℝ^*m*^, 𝔼 ∈ ℝ^*n*^, 𝕎∈ℝ^*n*^) for investigating the advection-diffusion-reaction dynamics. The cardinality of all the properties and the operators defined on 𝒢 are summarized in supplementary S1 Table.

**TABLE 1.** Specifications of the computational domain, values of flow boundary conditions, and the values of transport parameters used in the model.

| Tissue (unit) | Islet | Tumor Design 1 & 2 | Mesentery |
| --- | --- | --- | --- |
| Image dimensions |  |  |  |
| x (pixels) | 366 | 529 | - |
| y (pixels) | 366 | 529 |  |
| z (pixels) | 253 | 348 |  |
| Image spacing($\mu m$) | 0.6 | 2.31 | - |
| Domain size |  |  |  |
| x ($\mu m$) | 219.6 | 1221.9 | - |
| y ($\mu m$) | 219.6 | 1221.9 | |
| z ($\mu m$) | 151.8 | 803.8 | |
| # Vessel segments | 52 | 63 | 489 |
| $v_{in}$ ($\mu m/s$) | 160<sup>a</sup> | 400<sup>b</sup> | 400<sup>b</sup> |
| $Q_{in}/Q_{out}$(nl/min) | 3.76 | 23.8 | 40 |
| $P_{in}$ (mmHg) | 60 | 60 | - |
| $P_{out}$ (mmHg) | - | - | 0 |
| $\nu$ (Pas) | 0.004<sup>c</sup> | 0.004<sup>c</sup> | 0.004<sup>c</sup> |
| $\tilde{D}_A$ ($cm^2/min$) | 5.46e-4<sup>c</sup> | 5.46e-4<sup>c</sup> | 5.46e-4<sup>c</sup> |
| $\tilde{D}_B$ ($cm^2/min$) | 7.71e-4<sup>c</sup> | 7.71e-4<sup>c</sup> | 7.71e-4<sup>c</sup> |

**Figure 2.**
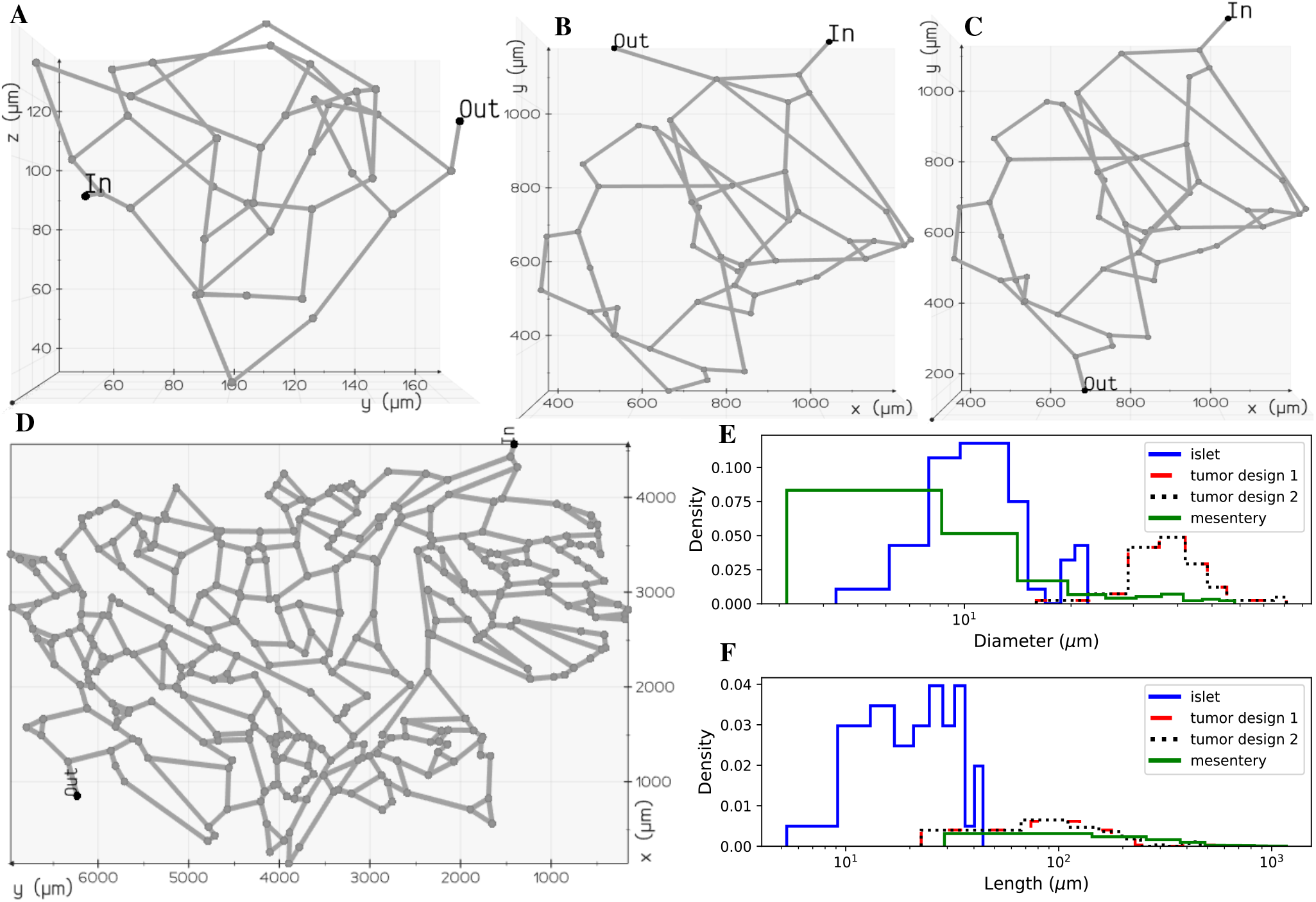
Microvascular network configurations: **(A)** pancreatic islet, **(B)** tumor design 1, **(C)** tumor design 2, and **(D)** mesentery generated after skeletonization of 3D volumes reconstructed from the image geometries studied in (Chen et al., 2020) and (d’Esposito et al., 2018). Diameter **(E)** and length **(F)** distributions of the vessel segments present in the four geometries.

### 2.2 Preliminary Assumptions

The time-dependent uptake and release of biochemicals by a tissue is determined by the gradient-driven transport across the capillary vessels which facilitate the transvascular exchange of solute with the tissue interstitium. Subsequently, the carrier-mediated sites present on the plasma membrane of cells aid the uptake of nutrient resources from the interstitial space for metabolism. Here we introduce a graph-based mathematical framework for capturing blood-tissue exchange. The following simplifying assumptions are made in our model similar to the assumptions considered in the other multiscale studies. (i) Assuming the scalar concentration in the interstitial fluid attains rapid-equilibrium with the concentration in the blood (Chalhoub et al., 2007), the interstitial compartment is not modeled. (ii) The endothelial layer of the capillary surface is lined with metabolite transporters which promote facilitated diffusion of biochemicals across the capillary wall, a similar approach has been presented in Heaton et al. (Heaton, 2012). We consider this as a reasonable assumption since a large fraction of the endocrine cells of the pancreas lie in close contact with the surface of the capillaries (Cohrs et al., 2017). (iii) Due to the deficiency of lymphatic drainage in the islets of Langerhans (Korsgren and Korsgren, 2016), fluid exchange with lymph vessels is not modeled (Thurber and Weissleder, 2011). Based on these considerations we subdivide each blood vessel branch and the layer of tissue surrounding the capillary into discrete functional units, diagrammed in Figure 3**D**. The molar transport occurring through the blood vessel compartment of these functional units is modeled by one dimensional advection-dispersion equation (Taylor, 1954) that accounts for convection flux, the axial and radial diffusive flux of the solute molecules. The transcapillary exchange flux and the cellular processes occurring in the tissue compartment of each functional unit are modeled by rate expressions that capture the kinetics of metabolite specific transporters and enzyme catalyzed reactions, respectively. The governing equations that model the inter-compartment dynamics of these well-mixed volumes embedded in the finite connected network representation of the capillary bed is detailed in Section 2.4 and 2.5.

**Figure 3.**
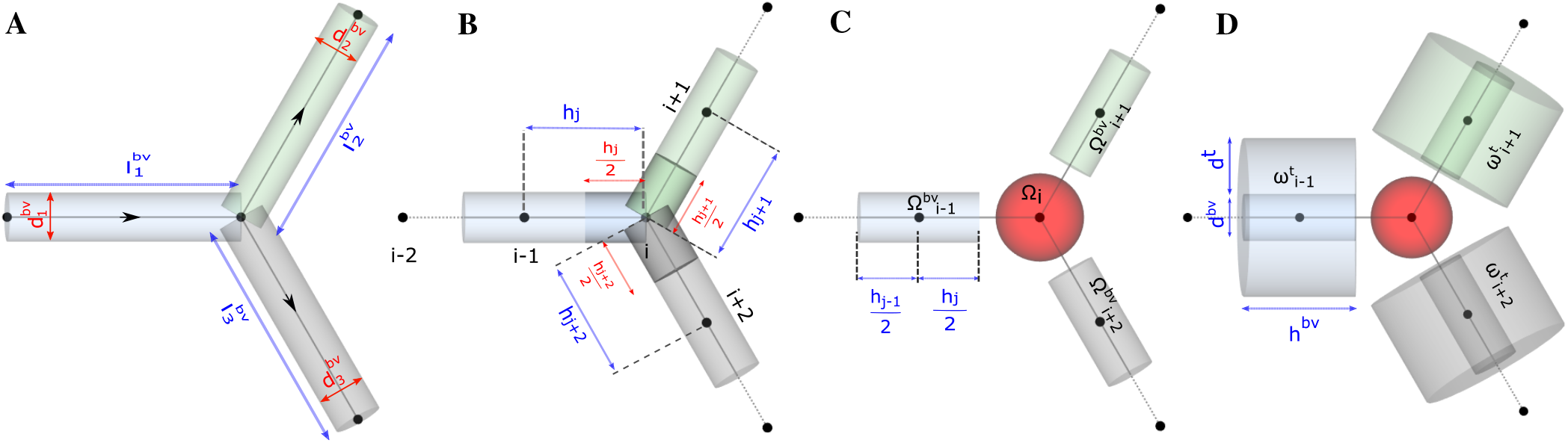
Schematic representation of functional units in the model. Computational domain for studying **(A)** flow distribution, **(B)** and **(C)** advective-dispersive transport of biochemicals in the discrete volumes of blood vessel Ω^*bv*^, **(D)** advective-dispersive-reactive transport of biochemicals in the discrete volumes of blood vessel Ω^*bv*^ and tissue domains *ω*^*t*^.

### 2.3 Mathematical Formulation of Flow Distribution in the Network

To simulate the spatio-temporal evolution of chemical species, we first solve for the flow field in the vascular branches using the approach generally applied in studies that focus on modeling flow distribution in branching networks (Erlich et al., 2019, d’Esposito et al., 2018, Kirkegaard and Sneppen, 2020, Poelma, 2017).

#### 2.3.1 Domain

To perform fluid flow simulations, the oriented graph depicted in Figure 3**A** was used as the computational domain for setting up the linear system of equations. As illustrated, each blood vessel branch was treated as an axially symmetric cylinder of axial length *l*^*bv*^ and circular radius *r*^*bv*^ derived from diameter *d*^*bv*^ extracted from the skeletonized geometry (Sec.2.1).

#### 2.3.2 Equations

We consider blood as an incompressible, viscous Newtonian fluid and apply Hagen-Poiseuille equation (Eq. 2) for modeling the conductance of an edge as a function of viscosity *μ*, radius *r*^*bv*^, length *l*^*bv*^. The linear analysis of the flow distribution presented here is applicable for laminar flow, *Re <* 1 in all segments and this can be extended further to study the nonlinear rheology of blood using the procedure illustrated in Pries et al. (Pries et al., 1990). Therefore, the distribution of flow in a microvascular network is determined by the pressure gradient and the resistance offered to flow.

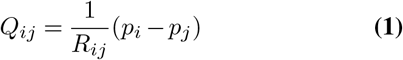

Here *p*_*i*_ and *p*_*j*_ are the pressures at tail *i* and head *j* nodes of the oriented edge *e*_*ij*_, *G*_*ij*_ is the conductance associated with *e*_*ij*_.

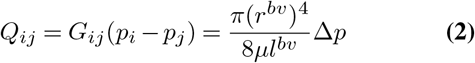

The vector *Q* ∈ ℝ^*n*^ of volumetric flow rates in *n* edges is defined in Eq. 3. The negative of the oriented incidence matrix *M* (𝒢) ∈ ℝ^*n*×*m*^, denoted as *M* henceforth, is the gradient operator that acts on the vector *P* ∈ ℝ^*m*^ of nodal pressures to result in the vector Δ*P* ∈ ℝ^*n*^ of pressure gradients. We obtain *Q* by premultiplying Δ*P* with the diagonal conductance-matrix *G* ∈ ℝ^*n*×*n*^, this scales the pressure gradient across each edge by the corresponding edge conductance.

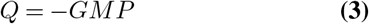

In our model, we consider blood as an incompressible fluid and determine the unknown nodal pressures by imposing mass conservation at all nodes. Consequently, the net flow at any given node *i* is zero (Eq. 4). Here, *Q*_*ij*_ is positive when flow enters node *i*(*Q*_*i*←*j*_), negative when flow leaves node *i*(*Q*_*i*→*j*_), and 𝒜(*i*) denotes the set of nodes that are adjacent to *i*.

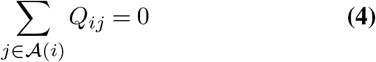

Over *m* nodes in the network, the vector *M*^*T*^ *Q* (*M*^*T*^ ∈ ℝ^*m*×*n*^ is divergence operator which is given by the transpose of *M*) defines the flow conservation at all nodes excluding the terminal nodes where the flow boundary conditions are specified. The non-zero entries of vector 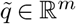 (Eq. 5) contains the values of inflow or outflow rates at the boundary nodes (Erlich et al., 2019). In addition to the flow rate, we specify one value of known pressure at the inlet *p*_*in*_ or outlet *p*_*out*_. The values of these boundary conditions were specified based on experimental measurements of blood flow velocities reported in Table 1.

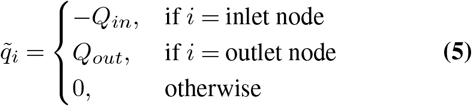

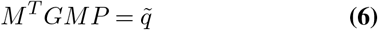

Here *M*^*T*^ *GM* is a square matrix. After substitution of known pressures in the vector P, the columns of *M*^*T*^ *GM* scaled by the values of known pressure are shifted to the RHS of Eq. 6. This operation results in a non-square matrix on the LHS of Eq. 6. The resulting system of linear equations is solved for the unknown nodal pressures by finding the pseudoinverse (Golub and Pereyra, 1973) which is the generalization of inverse for rank-deficient matrices. Pseudoinverse was computed in MATLAB using singular-value decomposition. From the estimated nodal pressures, the centerline velocity *u*_*ij*_ and the volumetric flow rates are computed using Eq. 2.

### 2.4 Advection-Dispersion of Chemical Species in Blood Vessel

#### 2.4.1 Domain discretization

The axial lines (edges) of the pipe network in Figure 3**A** was spatially discretized to set up the computational domain and study the transport of biochemical species in the microvasculature. For discretizing the edges into 1D elements, in Gmsh (Geuzaine and Remacle, 2009), the vasculature was represented as a geometry with point and line entities. The length of the mesh elements, denoted by *h* in Figure 3**B**, were constrained by specifying the maximum and minimum characteristic lengths i.e. *h* ∈ [*c*_*l*_ + *δ, c*_*l*_ -*δ*]. We derive the characteristic length *c*_*l*_ based on the average diameter of a biological cell, approximately 11.5 *μ*m, calculated from the volume ranges reported in pancreas (Pisania et al., 2010, Parween et al., 2016) and we consider a deviation *δ* of 2.5 *μ*m from *c*_*l*_.

#### 2.4.2 Domain volume elements

Each node in the discretized domain forms the center of the volume surrounding it. For instance, the volume of *i*^*th*^ node 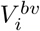 (Equation 7) located at the bifurcation point depicted in Figure 3**B** is the sum of half-cylinder volumes formed between nodes i-j where j ∈ 𝒜 (*i*) (Reichold et al., 2009). Here, 𝒜 (*i*) is the set {*i*-1, *i*+1, *i*+2} of nodes adjacent to *i, A*_*ij*_ and *l*_*ij*_ are the cross-sectional area and length of the cylindrical volume between nodes i-j, respectively. At the junction nodes, we merge the half cylinder volumes of the adjoining edges and equate the sum to a spherical volume (Figure 3**C**). Therefore, each branch in the network is dissected into cylindrical elements and the branches are assembled together by the spherical elements at the junctions. Since the rate of momentum transport is three orders of magnitude greater than the rate of mass transport, the accumulation term at any N-furcation junction is zero while solving for flow field and non-zero while solving the mass transport problem. As an example, for glucose species, the ratio of momentum to mass transport defined by Schmidt number *ν/D* is around 4000.

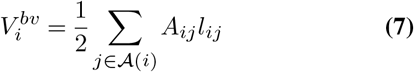

#### 2.4.3 Equations

The continuous formulation of the advection-dispersion physics is given by the following partial differential equation which describes the solute mole balance.

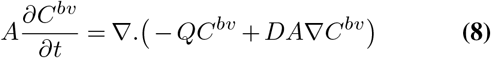

Here, *A* denotes the area of flow, *D* is the dispersion coefficient of species, Q denotes the flow field, and *C*^*bv*^ is the scalar concentration of a chemical species. To shift from the continuous to discrete counterpart, first we assign the scalar concentration field *C*_0_ to all vertices, and the flow field and dispersion coefficients form the edge weights. The net change in molar concentration of a species 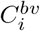 in the control volume 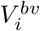 at the *i*^*th*^ node of the blood vessel depends on the contributions from advective 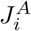 and dispersive fluxes 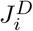 presented in equation 9. The *influx* and *outflux* of 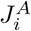 are the summation of molar fluxes through the edges feeding into *i*(*j → i*) and leaving *i* (*i*→*l*), respectively. And the molar flux through an edge is the flow-weighted concentration of the compartment from which the oriented edge originates (Chapman, 2015, Hošek and Volek, 2019). The third term contributes to the dispersive transport. Further, 𝒜 (*i*) is the set that contains nodes that are adjacent to node *i*, 𝒜 _+_(*i*) is the set of in-neighbor nodes of *i*, 𝒜 _−_(*i*) is the set of out-neighbor nodes of *i, D*_*ik*_, *A*_*ik*_ and *l*_*ik*_ are dispersion coefficient, cross-sectional area and length of edge *e*_*ik*_, respectively.

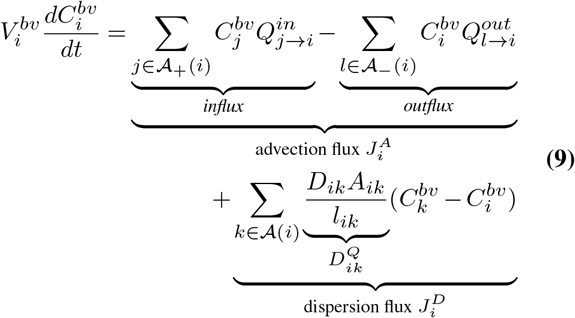

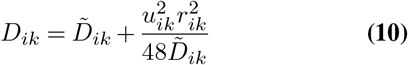

The dispersion coefficient is determined from the Aris-Taylor’s relation in Eq. 10. Here,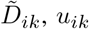, *u*_*ik*_ and *r*_*ik*_ are the diffusion coefficient of a species, centerline velocity and radius of *e*_*ik*_, respectively.

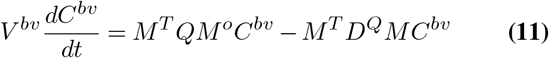

For setting up the discrete advection-diffusion equation on the network, we express the first and second order partial derivatives in Eq. 8 in terms of weighted advection Laplacian proposed by Rak et al. (Rak, 2017) and weighted diffusion Laplacian, respectively. This yields a system of ordinary differential equations shown in Equation 11, 12. In Eq. 11,*D*^*Q*^ ∈ ℝ^*n*×*n*^ is the diagonal matrix with diagonal entries the volumetric dispersion coefficient *DA/l* of each edge specified in Eq. 9, *Q* ∈ ℝ^*n*×*n*^ is the diagonal matrix with diagonal entries the volumetric flow rate of each edge, *M*^*o*^ ∈ ℝ^*n*×*m*^ is the modified incidence matrix (Erlich et al.,2019, Rak, 2017), *M*^*T*^ *D*^*Q*^*M* is the weighted dispersion Laplacian matrix *L*^*D*^ (𝒢) ∈ ℝ^*m*×*m*^ and *M*^*T*^ *QM* ^*o*^ is the weighted advection Laplacian matrix *L*^*Q*^ (𝒢)∈ℝ^*m*×*m*^ (Eq. 12).

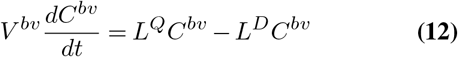

### 2.5 Advection-Dispersion-Reaction

#### 2.5.1 Domain

We adapt the Krogh cylinder approach presented in multi-scale models (Chalhoub et al., 2007, Berndt et al., 2018b, Thurber and Weissleder, 2011) and approximate the layer of tissue surrounding the vessels as a hollow cylindrical volume element *ω*^*t*^ illustrated in Figure 3**D**. The outer diameter of the region *ω*^*t*^ is given by the summation of *d*_*bv*_ and *d*_*t*_. A value of 12.4μm was used in our model for *d*_*t*_. This value is derived from the physical volume of an insulin secreting beta cell (1020*μm*^3^) (Finegood et al., 1995) located in the islets of pancreas.

#### 2.5.2 Equations

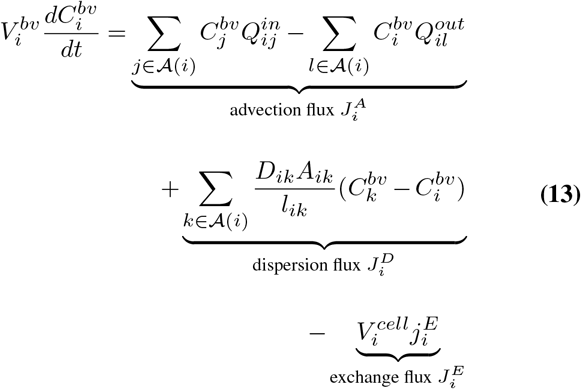

In the last term of Eq. 13, 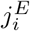 (Eq. 15) is the net exchange rate that governs the bidirectional transport between tissue compartment 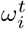 and blood vessel compartment 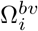 of the functional unit. The uptake or release flux 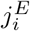 is computed by multiplying 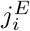 with 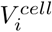 since the maximal rates are often reported in per unit volume of a biological cell. This ensures mole balance when a species moves from 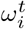 to 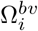 which differ in volumes. Further, 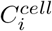 is the concentration of species in the tissue cell that interacts with *i*^*th*^ node of Ω^*bv*^, the half-saturation constant *K*_*m*_ quantifies the affinity of transporter-protein or an enzyme for a metabolite, and *V*_*m*_ is the maximal rate of metabolite transport. When 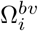 is encompassed by 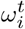, interaction between the node associated with both the compartments exists and 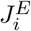 appears as a source or sink term in Eq. 13. At the junction nodes of blood vessel where no interaction exists 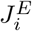 is zero.

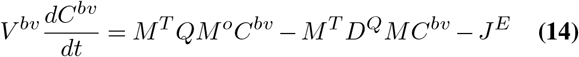

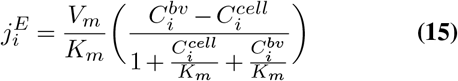

As an example, we consider a minimal model of glucose metabolism in *β*-cells in the islets. At basal condition, the concentration of glucose and lactate are at basal level in the bloodstream. In fed state, glucose transporters sense the high blood glucose level and export glucose from Ω^*bv*^ to Ω^*t*^. The glucose-to-lactate conversion in Ω^*t*^ is presented as a lumped reaction in our model for simplicity. Ω^*t*^ acts as a source of lactate and the lactate transporter facilitates its release into the bloodstream. The direction of the exchange flux is dictated by the concentration gradient *C*^*bv*^ −*C*^*cell*^ across the vessel wall.

### 2.6 Cell

The mole balance of each species in the cell domain *ω*^*t*^ is modeled by

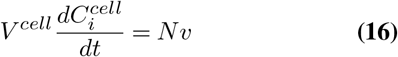

where, N is the stoichiometric matrix and *v* is the reaction flux vector. The rate expressions of the kinetic reactions and metabolite transporters, and the values of the kinetic parameters used in the model are presented in Table 2. The values of half-saturation constants, *K*_*m*_ and *V*_*m*_ were chosen from *panmin* (König and Deepa Maheshvare, 2021), a minimal model of glucose metabolism and insulin secretion in the pancreatic *β*-cell. The *V*_*m*_ values from the minimal model were scaled in this study so that the reaction fluxes are comparable to the diffusive and convective fluxes. To apply our framework for predictive modeling in clinical applications, the *V*_*m*_ values can be determined by model calibration to achieve good agreement between model predictions and experimental measurements. Steady-state and transient values of metabolite and flux distributions measured from biochemical assays can be used as the inputs for calibrating the tunable parameters in the model (Berndt et al., 2018b). Since the focus of this study is in introducing and illustrating the applicability of our mathematical framework for bridging multiple scales, the parameters were not calibrated in the glucose-lactate test system presented here.

**TABLE 2.** Reactions, rate laws and parameter values of exchange and cellular reactions.

| Flux | Reaction | Rate law | Parameter values ( $V_m$ in M/min, $K_m$ in mM ) |
| --- | --- | --- | --- |
| $v_{TI}$ | $A^{bv} \rightleftharpoons A^{cell}$ | $\frac{V_m^{T1}(A^{bv} - A^{cell})}{K_{m,A}^{T1} + A^{bv} + A^{cell}}$ | $V_m^{T1} = 10, K_{m,A}^{T1} = 1.0$ |
| $v_E$ | $A^{cell} \rightarrow 2B^{cell}$ | $\frac{V_m^E A^{cell}}{A^{cell} + K_{m,A}^E}$ | $V_m^E = 0.01, K_{m,A}^E = 4.5$ |
| $v_{T2}$ | $B^{cell} \rightleftharpoons B^{bv}$ | $\frac{V_m^{T2}(B^{cell} - B^{bv})}{K_{m,B}^{T2} + B^{cell} + B^{bv}}$ | $V_m^{T2} = 10, K_{m,B}^{T2} = 0.5$ |

A: glucose, B: lactate, T1: glucose transporter *glcim*, T2: lactate transporter *lacex*, E: glucose to lactate converter *glc2lac* are the tags used in the mathematical model.

## 3 RESULTS AND DISCUSSION

The discrete model framework developed allows to bridge the cell-to-vessel exchange and explicitly model the cellular dynamics. In this section, the mathematical formulations presented in Section 2 are verified using several test systems and the results are compared with finite element analysis in COMSOL. The analysis includes solving two steps that are coupled: first, a flow field analysis which involves computation of pressure and velocity distribution in the blood vessel branches; second, analysis of the spatio-temporal evolution of the concentration fields.

We first present the results of pressure gradient and flow distribution observed across the islet vasculature. After validating the results of nodal pressures and edge velocities with the results from COMSOL, we proceed with the simulations of advection-dispersion dynamics of glucose species in the blood vessel. Here we compare the transient change in the blood glucose concentration obtained from our discrete model versus COMSOL simulations for islet and mesentery networks. Next, we investigate the effect of change in perfusion pattern on glucose distribution by varying the inlet and outlet locations in the tumor vasculature. Further, we examine the effect of different pressure gradients applied across the network and the effect of glucose doses supplied at the inlet on metabolite rise times observed in the islet vasculature. All the 3D visualizations of flow and concentration fields presented in this paper are rendered using *vedo* (Musy et al., 2021), a python based module for analyzing and visualizing multi-dimensional point-cloud, mesh and volume data.

### 3.1 Comparison of Flow and Concentration Fields

To illustrate how the results from our discrete formulation compare with the finite element implementation available in COMSOL, we first solve the static flow problem (Eq. 6) and use the flow profile for simulating the advection-dispersion dynamics (Eq. 12) in the islet vasculature. The pressure and velocity field computed across the islet vasculature are shown in Figure 4**A**,**B**. We observe a net pressure drop of 34.83Pa for an inlet pressure and flow rate of 60Pa and 3.76nl/min, respectively. The velocity distribution lies in the range reported by Diez et al. (Diez et al., 2017). The conservation of flow at each node is cross-verified by computing the divergence of the flow field *M*^*T*^ *Q* and the visualization is presented in Figure 4**C**.

**Figure 4.**
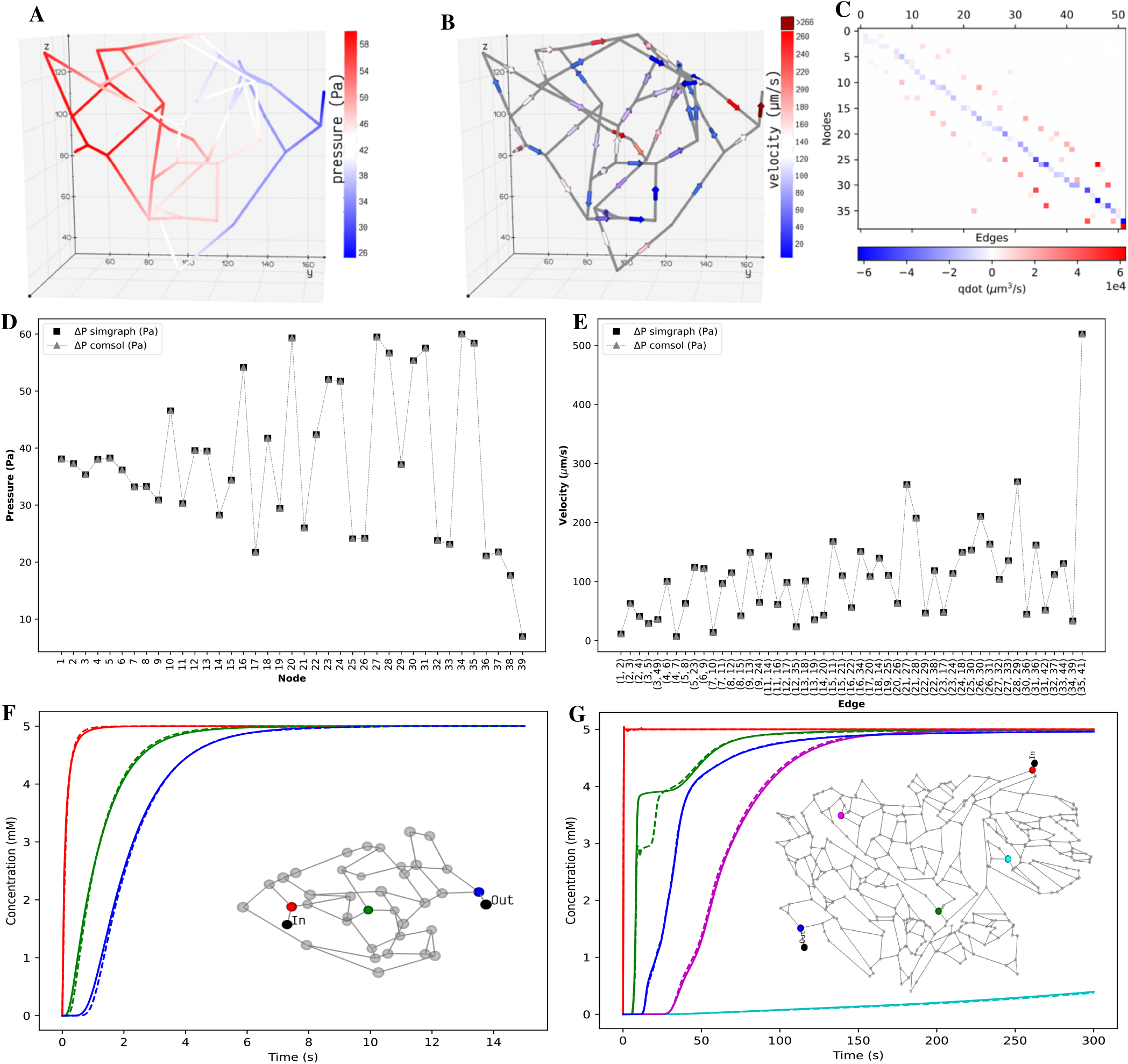
Distribution of the static properties of islet vasculature. **(A)** Pressure gradient from inlet to outlet and **(B)** directional flow from inlet to outlet. **(C)** Flow conservation at all nodes. The summation of flows through the edges entering and leaving each node, except the boundary nodes, add to zero. **Validation of static results of the islet vasculature. (D)**: Comparison of nodal pressures between the current work and COMSOL simulation. **(E)**: Comparison of edge velocities between the current work and COMSOL simulation. **Comparison of the concentration profiles at various locations in the islet and mesentery vasculatures. (F)** Scalar concentrations observed at positions 12.07 *μ*m (red), 54.85 *μ*m (green), and 110.27 *μ*m (blue) from the inlet node. **(G)** Scalar concentrations observed at positions 1267.78 *μ*m (blue), 1692.3 *μ*m (cyan), 3476.96 *μ*m (green), 5085.33 *μ*m (magenta), and 6319.57 *μ*m (blue) from the inlet node. Solid and dashed lines indicate the transient change in glucose concentration from our model and COMSOL, respectively.

#### 3.1.1 Validation in COMSOL Multiphysics

Here we describe the simulation implementation of the coupled multiphysics problem in COMSOL to validate the flow fields and concentration fields of advection-dispersion simulations from our discrete model. *Geometry:* The generation of the geometrical model of the islet and mesentery vasculature (Figure 2**A**,**D**) was automated using an AutoLISP script (Mac, 2020). The coordinates of points and the connectivity information of the lines were specified as inputs for creating the CAD geometry. The DXF file containing the geometry data was imported in COMSOL and *Normal* mesh size was used to generate the mesh elements. *Parameters:* Values of diffusion coefficient of glucose species, viscosity of blood, inlet pressure and outlet flow rate specified in Table 1, and diameter of all branches in the vasculature were defined as input parameters. *Computation*: Fluid flow was studied as static problem in the *Pipe Flow* module considering blood as a Newtonian fluid. This stationary problem was solved as a linear system using the *Direct* solver by specifying the pressure and flow boundary conditions. Then, one-way coupling of the flow physics was established with *Transport of Dilute Species in Pipes* module to solve for the time-dependent advection-dispersion physics of glucose species in the finite element solver. For this transient simulation, a value of 5mM was used for the Dirichlet boundary condition defined at the inlet and mass outflow was modeled by setting the diffusive flux to zero. The initial concentration was set to zero in the volume elements of our discrete framework. In COMSOL, the concentration was initialized to zero and a smoothed step function was applied to avoid discontinuity with the boundary condition.

Figure 4**D**,**E** show the nodal pressures and edge velocities computed from our model are consistent with the results from COMSOL for the islet vasculature. Time-varying concentration profile of glucose species obtained from our discrete model is compared with COMSOL simulations at nodes highlighted in the inset of Figure 4**F**. After comparing the results of the islet vasculature containing 52 edges and 125 discretized elements in our discrete model, we further extend this analysis to compare the results (Figure 4**G**) of the large mesentery network with 489 edges and 9033 discretized elements. Supplementary videos S1 and S2 show the comparison of evolution of concentration profiles from our model versus the results from COMSOL for the entire region of the islet and mesentery vasculature, respectively.

#### 3.1.2 Scalability

The procedure involved in solving the partial differential equations can be split into two steps: assembly and solve step. In the assembly step, discretization is done and matrices are generated for the second step which solves the system. For the islet vasculature, we report the time taken for solving the stationary fluid flow and the transient advection-dispersion dynamics for a time span of 15s. Our discrete method implementation in MATLAB takes 7s and the finite element solver in COMSOL takes 10s. For the mesentery network, running the advection-dispersion dynamics for a time span of 300s takes 267s and 168s in MATLAB and COMSOL implementations, respectively. We provide the sparse Jacobian pattern as an additional input to *ode15s* to speed up the compute time. Solving the same system in Julia using QNDF method, which is a translation of the MATLAB’s ode15s, gives a speed up of 127x when compared to COMSOL. A relative error tolerance of 1e-3 is set for carrying out all simulations and an absolute error tolerance of 1e-6 is set for both ode15s and QNDF.

Some of the challenges involved in expanding this approach to a vascular network composed of arterioles, arteries, and veins would be in scaling flow parameters and resistances to flow (such as molecular interaction in capillary versus viscosity in larger vessels) to fit the formulation so that the model still retains its physical fidelity. Computationally, there can be stiffness in the differential equations and condition numbers of matrices may be affected, especially when both capillary and larger vessels are present in the model. This may need smaller time steps in simulation and regularization of affected matrices.

### 3.2 Influence of Flow Topology on Scalar Transport

Engineered perfusable vasculatures have been useful for investigating the structure-function relationship of complex vasculatures (Kinstlinger et al., 2020). Computational models that capture the influence of flow topology on the scalar transport will enable experimental scientists to design and test the efficacy of optimized drug delivery systems.

Motivated by the experiments carried out by Chen et al. (Chen et al., 2020) on microfluidic devices imprinted with blood vessel vasculatures, we study the effect of variation in perfusion patterns on the metabolite rise times *t*_*r*_ in two different geometric configurations (Figure 2**B**,**C**) of the tumor vasculature examined in their study. Flow distribution was computed for an inlet pressure of 60Pa and fluid flow rate of 23.8nl/min. Spatial distribution of glucose species was obtained and the rise time in each volume element 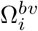 is computed by finding the time taken for the concentration to rise from 10% to 90% of the steady-state value. Figure 5 illustrates the time taken for the distribution of glucose in *Tumor Design 1* (Figure 5**B**) is much greater when compared to *Tumor Design 2* (Figure 5**A**). These findings support the dye distribution patterns observed in these two configurations reported in Chen et al.. Supplementary video S3 shows the distribution of glucose computed by our discrete model for both the configurations of tumor vasculature.

**Figure 5.**
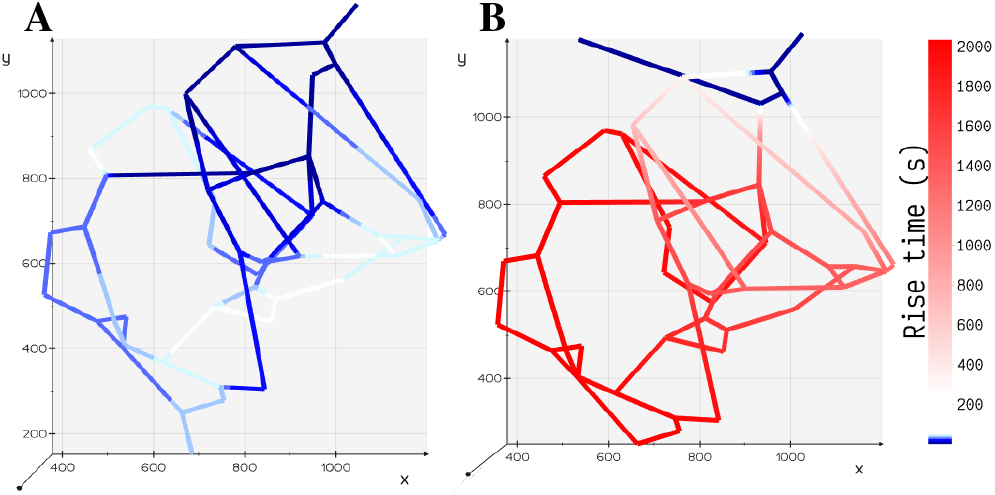
Comparison of rise time *t*_*r*_ for two different configurations of tumor vasculature. **(A)** Tumor Design 2: the outlet is positioned away from the inlet and the rise time is shorter in this configuration. **(B)** Tumor Design 1: the outlet is positioned closer to the inlet and the rise time is longer in this configuration.

### 3.3 Functional Coupling of Blood Vessel-Cell Exchange

In the mathematical framework presented in Section 2, it is practical to model heterogeneous cell types exhibiting heterogeneous enzyme activity at different vascular and cell densities observed in various pathophysiological conditions.

In this study, for the ease of demonstration of our method, we present the results of a minimal model of cellular glucose metabolism with uniform enzyme activity and homogeneous cell type exchanging biochemicals with bloodstream. In Figure 6, we present the volumetric spatial analysis of the islet vasculature enveloped by a layer of homogeneous cell mass which forms the annular region of the tissue domain *ω*^*t*^. The extracellular glucose and lactate concentrations in Ω^*bv*^ are initialized to 5mM and 1.2mM at the network inlet. The intracellular concentrations of glucose and lactate in *ω*^*t*^ are initialized to zero. Due to the high concentration of glucose in the bloodstream, the gradient established between Ω^*bv*^ and *ω*^*t*^ in response to the advective-dispersive transport promotes the uptake of glucose by the annular region *ω*^*t*^. The cellular enzymes metabolize glucose-to-lactate and the concentration of lactate in *ω*^*t*^ rises above the basal value in the bloodstream. This gradient drives the export of lactate into the bloodstream until the system equilibrates.

**Figure 6.**
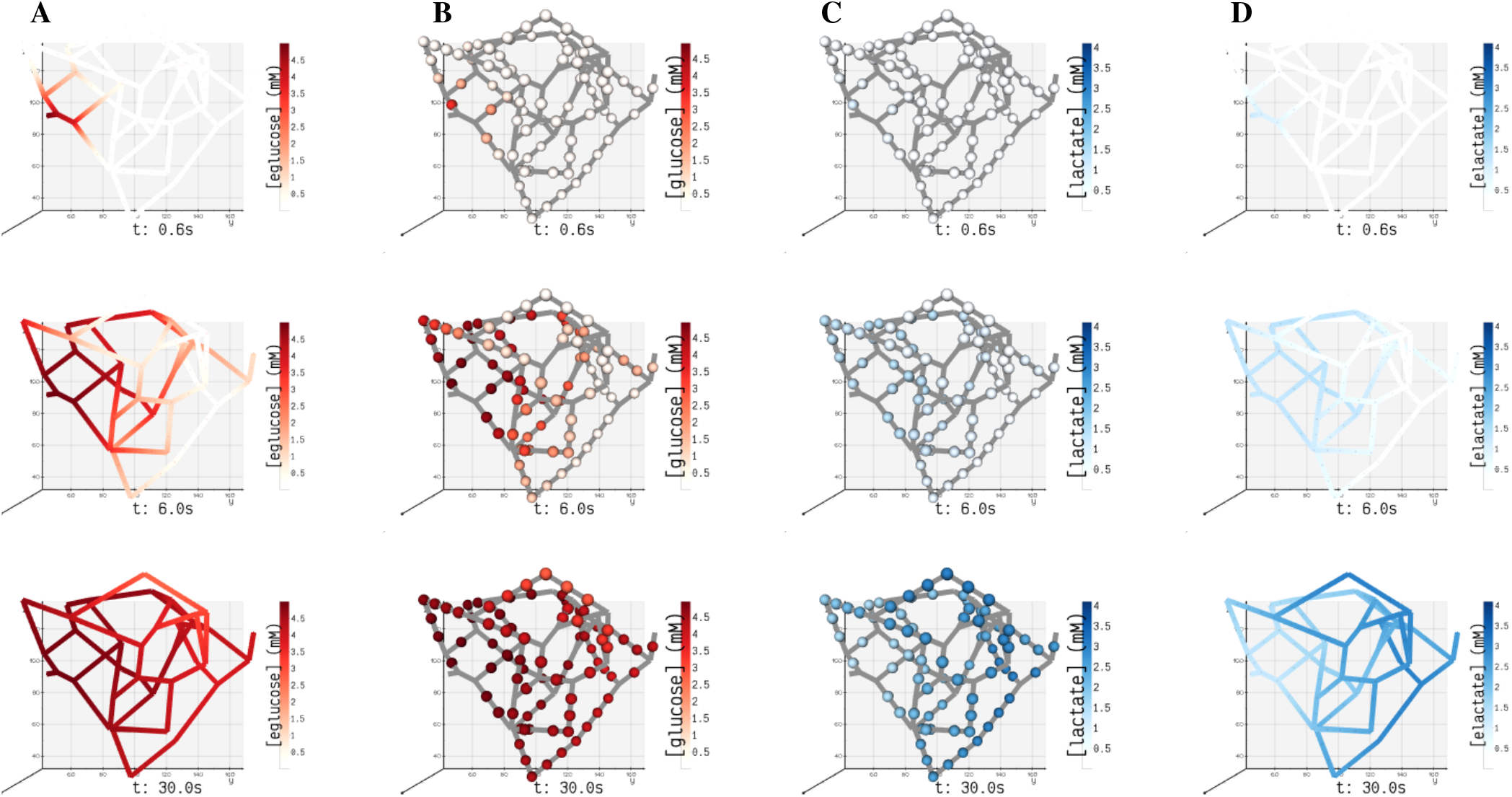
Spatial concentrations of metabolites in blood vessel and tissue domains of the islet vasculature observed at three different instants in time (0.6, 6, 30 seconds). **(A)** [eglucose]/ B [elactate] and **(C)** [glucose]/ **(D)** [lactate] denote the concentration of glucose/ lactate species observed in blood vessel Ω^*bv*^ and tissue *ω*^*t*^ compartments, respectively. In panel **(A)** and **(D)**, the concentration of cylindrical and spherical volume elements embedded in the blood vessel domain is simulated and the gradient displayed along the length of the blood vessel is generated by interpolation. In panel **(B)** and **(C)**, the color-coded spheres represent the concentration evaluated in the annular region of *ω*^*t*^.

It is known from clinical observations that vascular phenotypes alter in cohorts with disease conditions such as diabetes and tumor. For example, in a tumor condition the glycolytic enzymes are upregulated to produce more energy molecules that aid in the rapid proliferation of tumor cells (St Clair et al., 2018, Rojas et al., 2018). As a result, glucose-lactate dynamics is perturbed when compared to normal cells. In case of diabetes, histopathological and image reconstruction studies reveal that a decrease in the cell mass, reduction in vascular density can alter insulin release patterns (Cohrs et al., 2017, Richards et al., 2010). The framework proposed here can be useful in such clinical applications for carrying out systematic analysis where (i) the change in vascular density can be induced in the form of mutations in the network (e.g. by deleting or inserting blood vessels) to examine the effect of anatomical changes on functional response of a tissue; (ii) the alterations in the expression levels of enzymes quantified as fold changes in proteomics studies (Haythorne et al., 2019, Malinowski et al., 2020) can be mapped to the reaction velocities (i.e. Vm, a function of enzyme abundance, can be scaled using fold change) to examine the effect of genetic or environmental perturbations on cellular dynamics.

### 3.4 Sensitivity of Concentration Dynamics to ΔP and Glucose Dose

The pressure conditions observed in a vascular tissue may vary due to several physiological factors. To study the influence of pressure gradient on the time taken to reach the steady-state concentration, we vary ΔP across the vasculature by specifying the inlet pressure and zero outlet pressure. Figure 7**A** illustrates the concentration profiles generated by simulating the advection-dispersion dynamics of glucose species by varying the pressure drop from 20Pa to 200Pa. When ΔP is high, the velocity of fluid is high in each branch. Consequently, convective flow dominates over dispersion and this results in short rise times. The effect of change in pressure gradient results in change in transit time of the fluid from 90.76s at ΔP = 20Pa to 9.08s at ΔP = 200Pa. In Figure 7**B**, we show the sensitivity of the net glucose uptake flux and net lactate release flux to different glucose doses set at the inlet. It is observed that with increase in glucose dose the tissue units uptake more glucose from blood and this drives the formation of lactate in the tissue subdomains. The excess lactate is then transported to the vessel subdomains until equilibrium is attained and the driving potential is zero.

**Figure 7.**
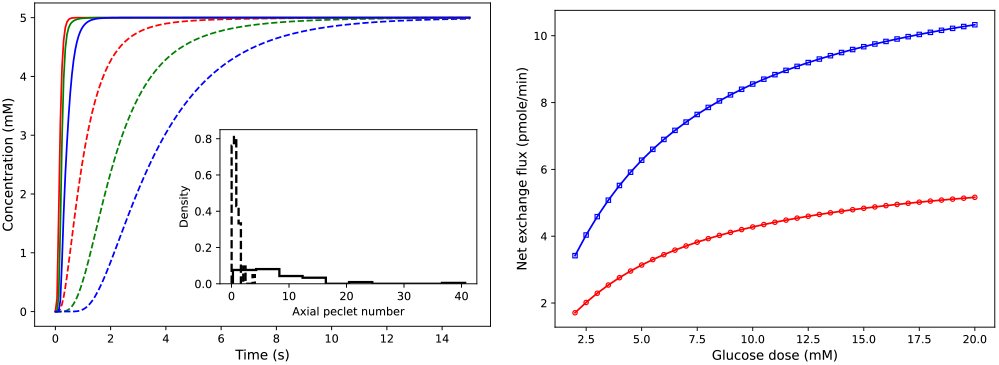
Sensitivity of concentration dynamics to ΔP and glucose dose. **(A)** Left: influence of varying pressure gradient across the network on metabolite rise times. Glucose concentration observed at positions 12.07 *μ*m (red), 54.85 *μ*m (green), and 110.27 *μ*m (blue) from the inlet node. Inset displays the distribution of peclet number obtained by computing 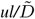 for each branch. Dashed and solid lines indicate the dynamic change in concentration observed at 20Pa and 200Pa, respectively. **(B)** Right: variation in the net glucose uptake (red) and net lactate release (blue) from the cells in response to change in glucose dose is displayed.

In conclusion, here we have presented a mathematical framework for understanding the multiscale connectivity existing in the functional networks of tissues. The test cases presented above demonstrate how experimental data from different sources (e.g. kinetic data available in databases such as SABIO-RK (Wittig et al., 2012) and BRENDA (Chang et al., 2021), proteomics and metabolomics data, imaging data of vascular phenotypes, flux measurements from perfusion experiments) can be encoded in our framework to build explicit models of cellular and vessel-to-cell interaction dynamics to better predict pathomechanisms. At the intra-organ scale, this work can be further extended to include cell-to-cell communication networks and decipher the order of communication occurring in the microenvironments with different vasculature architectures (e.g. periphery to the center, center to periphery, one pole to other patterns in islets (ElGohary and Gittes, 2018)) and cytoarchitectures (e.g. mantle-core and heterogeneous distribution of *β* and *α* cells in islets (Dolenšek et al., 2015)). The metamodel is easily mutable; it is possible to induce different vascular phenotypes and disease states by altering vascular density and incorporating fold changes of metabolic and enzyme concentrations in the cellular units. To further research efforts involved in carrying out virtual experiments of inter-organ communication, our framework can be utilized for piecing together the top-down and bottom-up modeling approaches and accommodate each organ at the desired spatial resolution. For example, a comprehensive understanding of multiorgan disease states such as diabetes can be developed by interpreting the intra- and inter-organ interaction as communication occurring within “network of networks”; in whole body models, the inter-organ communication can be modeled by abstracting organs as compartments forming nodes of the global network and the detailed local dynamics occurring in the functional networks of an organ can be modeled by including subnetworks in the global network.

## DATA AND SOFTWARE AVAILABILITY

All model simulations were performed with stiff-equation solver in MATLAB, Release R2019b, The MathWorks, Inc., Natick, Massachusetts, United States. Code is available upon request.

## ACKNOWLEDGMENTS

DM thanks Dr. Matthias König, Institute for Theoretical Biology, Humboldt-University Berlin for the valuable discussions, and Dr. Pradip Dutta, Department of Mechanical Engineering, Indian Institute of Science for offering the computational facility to carry out COMSOL simulations. Funding: Research of DM is supported by the Senior Research Fellowship from the Ministry of Human Resource Development (MHRD), Government of India. The preprint of this article has appeared online on bioRxiv (Deepa Maheshvare et al., 2021).

## AUTHOR CONTRIBUTIONS

DM, SR and DP conceived and designed the study. DM implemented the code, performed the analysis and drafted the manuscript. All authors read, discussed the results, revised and approved the manuscript.

## SUPPLEMENTARY MATERIAL

*Supplementary Material 1*

## REFERENCES

Matthias König, Sascha Bulik, and Hermann-Georg Holzhütter. Quantifying the contribution of the liver to glucose homeostasis: a detailed kinetic model of human hepatic glucose metabolism. PLoS Comput Biol, 8(6):e1002577, 2012.

Nikolaus Berndt, Sascha Bulik, Iwona Wallach, Tilo Wünsch, Matthias König, Martin Stockmann, David Meierhofer, and Hermann-Georg Holzhütter. Hepatokin1 is a biochemistry-based model of liver metabolism for applications in medicine and pharmacology. Nature communications, 9(1):1–12, 2018a.

Maria Masid, Meric Ataman, and Vassily Hatzimanikatis. Analysis of human metabolism by reducing the complexity of the genome-scale models using redhuman. Nature communications, 11(1):1–12, 2020.

Vishnu P Rao and Megan A Rizzo. Diffusion of metabolites across gap junctions mediates metabolic coordination of β-islet cells. bioRxiv, 2020.

Margaret Watts, Joon Ha, Ofer Kimchi, and Arthur Sherman. Paracrine regulation of glucagon secretion: the β/α/δ model. American Journal of Physiology-Endocrinology and Metabolism, 310(8):E597–E611, 2016.

Duk-Su Koh, Jung-Hwa Cho, and Liangyi Chen. Paracrine interactions within islets of langerhans. Journal of molecular neuroscience, 48(2):429–440, 2012.

Morten Gram Pedersen, Richard Bertram, and Arthur Sherman. Intra-and inter-islet synchronization of metabolically driven insulin secretion. Biophysical journal, 89(1):107–119, 2005.

Amlan K Barua and Pranay Goel. Isles within islets: the lattice origin of small-world networks in pancreatic tissues. Physica D: Nonlinear Phenomena, 315:49–57, 2016.

John Thomas Sorensen. A physiologic model of glucose metabolism in man and its use to design and assess improved insulin therapies for diabetes. PhD thesis, Massachusetts Institute of Technology, 1985.

MR Gray and YK Tam. The series-compartment model for hepatic elimination. Drug metabolism and disposition, 15(1):27–31, 1987.

Andreas Deussen and James B Bassingthwaighte. Modeling [15o] oxygen tracer data for estimating oxygen consumption. American Journal of Physiology-Heart and Circulatory Physiology, 270(3):H1115–H1130, 1996.

Elie Chalhoub, L Xie, V Balasubramanian, J Kim, and J Belovich. A distributed model of carbohydrate transport and metabolism in the liver during rest and high-intensity exercise. Annals of Biomedical Engineering, 35(3):474–491, 2007.

Jurij Dolenšek, Marjan Slak Rupnik, and Andraž Stožer. Structural similarities and differences between the human and the mouse pancreas. Islets, 7(1):e1024405, 2015.

Daniel A Beard and James B Bassingthwaighte. Advection and diffusion of substances in biological tissues with complex vascular networks. Annals of biomedical engineering, 28(3):253–268, 2000.

Qianqian Fang, Sava Sakadžić, Lana Ruvinskaya, Anna Devor, Anders M Dale, and David A Boas. Oxygen advection and diffusion in a three dimensional vascular anatomical network. Optics express, 16(22):17530, 2008.

Luke LM Heaton, Eduardo López, Philip K Maini, Mark D Fricker, and Nick S Jones. Advection, diffusion, and delivery over a network. Physical Review E, 86(2):021905, 2012.

M Kojic, M Milosevic, V Simic, EJ Koay, JB Fleming, S Nizzero, N Kojic, A Ziemys, and M Ferrari. A composite smeared finite element for mass transport in capillary systems and biological tissue. Computer methods in applied mechanics and engineering, 324:413–437, 2017.

Soroush Safaei, Pablo J Blanco, Lucas O Müller, Leif R Hellevik, and Peter J Hunter. Bond graph model of cerebral circulation: toward clinically feasible systemic blood flow simulations. Frontiers in physiology, 9:148, 2018.

Alexander Erlich, Philip Pearce, Romina Plitman Mayo, Oliver E Jensen, and Igor L Chernyavsky. Physical and geometric determinants of transport in fetoplacental microvascular networks. Science advances, 5(4):eaav6326, 2019.

Kaito Ii, Kota Mashimo, Mitsunori Ozeki, Takahiro G Yamada, Noriko Hiroi, and Akira Funahashi. Xitosbml: A modeling tool for creating spatial systems biology markup language models from microscopic images. Frontiers in genetics, 10:1027, 2019.

Nikolaus Berndt, Marius Stefan Horger, Sascha Bulik, and Hermann-Georg Holzhütter. A multiscale modelling approach to assess the impact of metabolic zonation and microperfusion on the hepatic carbohydrate metabolism. PLoS computational biology, 14(2):e1006005, 2018b.

Leonardo Bellocchi and Nikolas Geroliminis. Unraveling reaction-diffusion-like dynamics in urban congestion propagation: Insights from a large-scale road network. Scientific reports, 10(1):1–11, 2020.

Dinesh Kumar, Jatin Gupta, and Soumyendu Raha. Partitioning a reaction–diffusion ecological network for dynamic stability. Proceedings of the Royal Society A, 475(2223):20180524, 2019.

Christophe Besse and Grégory Faye. Dynamics of epidemic spreading on connected graphs. Journal of Mathematical Biology, 82(6):1–52, 2021.

Timothy S Frost, Victor Estrada, Linan Jiang, and Yitshak Zohar. Convection–diffusion molecular transport in a microfluidic bilayer device with a porous membrane. Microfluidics and Nanofluidics, 23(10):1–13, 2019.

Jianchen Yang, Tessa Davis, Anum S Kazerouni, Yuan-I Chen, Meghan J Bloom, Hsin-Chih Yeh, Thomas E Yankeelov, and John Virostko. Longitudinal fret imaging of glucose and lactate dynamics and response to therapy in breast cancer cells. Molecular imaging and biology, pages 1–12, 2021.

Nan Jiang, Roger D Cox, and John M Hancock. A kinetic core model of the glucose-stimulated insulin secretion network of pancreatic β cells. Mammalian Genome, 18(6):508–520, 2007.

Marc Prentki, Franz M Matschinsky, and SR Murthy Madiraju. Metabolic signaling in fuel-induced insulin secretion. Cell metabolism, 18(2):162–185, 2013.

Yih Yang Chen, Benjamin R Kingston, and Warren CW Chan. Transcribing in vivo blood vessel networks into in vitro perfusable microfluidic devices. Advanced Materials Technologies, 5(6):2000103, 2020.

Angela d’Esposito, Paul W Sweeney, Morium Ali, Magdy Saleh, Rajiv Ramasawmy, Thomas A Roberts, Giulia Agliardi, Adrien Desjardins, Mark F Lythgoe, R Barbara Pedley, et al. Computational fluid dynamics with imaging of cleared tissue and of in vivo perfusion predicts drug uptake and treatment responses in tumours. Nature Biomedical Engineering, 2(10):773–787, 2018.

Christoph Sommer, Christoph Straehle, Ullrich Köthe, and Fred A. Hamprecht. Ilastik: Interactive learning and segmentation toolkit. In 2011 IEEE International Symposium on Biomedical Imaging: From Nano to Macro, pages 230–233, 2011. doi: 10.1109/ISBI.2011.5872394.

Andriy Fedorov, Reinhard Beichel, Jayashree Kalpathy-Cramer, Julien Finet, Jean-Christophe Fillion-Robin, Sonia Pujol, Christian Bauer, Dominique Jennings, Fiona Fennessy, Milan Sonka, et al. 3d slicer as an image computing platform for the quantitative imaging network. Magnetic resonance imaging, 30(9):1323–1341, 2012.

Luca Antiga, Marina Piccinelli, Lorenzo Botti, Bogdan Ene-Iordache, Andrea Remuzzi, and David A Steinman. An image-based modeling framework for patient-specific computational hemodynamics. Medical & biological engineering & computing, 46(11):1097–1112, 2008.

Juan A Diez, Rafael Arrojo e Drigo, Xiaofeng Zheng, Olga V Stelmashenko, Minni Chua, Rayner Rodriguez-Diaz, Masahiro Fukuda, Martin Köhler, Ingo Leibiger, Sai Bo Bo Tun, et al. Pancreatic islet blood flow dynamics in primates. Cell reports, 20(6):1490–1501, 2017.

Emmanuel M Gabriel, Minhyung Kim, Daniel T Fisher, Colin Powers, Kristopher Attwood, Sanjay P Bagaria, Keith L Knutson, and Joseph J Skitzki. Dynamic control of tumor vasculature improves antitumor responses in a regional model of melanoma. Scientific reports, 10(1):1–13, 2020.

Luke L. M. Heaton. Biological transport networks. PhD thesis, University of Oxford, 2012.

Christian M Cohrs, Chunguang Chen, Stephan R Jahn, Julia Stertmann, Helena Chmelova, Jürgen Weitz, Andrea Bähr, Nikolai Klymiuk, Anja Steffen, Barbara Ludwig, et al. Vessel network architecture of adult human islets promotes distinct cell-cell interactions in situ and is altered after transplantation. Endocrinology, 158(5):1373–1385, 2017.

Erik Korsgren and Olle Korsgren. An apparent deficiency of lymphatic capillaries in the islets of langerhans in the human pancreas. Diabetes, 65(4):1004–1008, 2016.

Greg Michael Thurber and Ralph Weissleder. A systems approach for tumor pharmacokinetics. PloS one, 6(9):e24696, 2011.

Geoffrey Ingram Taylor. The dispersion of matter in turbulent flow through a pipe. Proceedings of the Royal Society of London. Series A. Mathematical and Physical Sciences, 223(1155):446–468, 1954.

Julius B Kirkegaard and Kim Sneppen. Optimal transport flows for distributed production networks. Physical Review Letters, 124(20):208101, 2020.

Christian Poelma. Exploring the potential of blood flow network data. Meccanica, 52(3):489–502, 2017.

Axel R Pries, Timothy W Secomb, P Gaehtgens, and JF Gross. Blood flow in microvascular networks. experiments and simulation. Circulation research, 67(4):826–834, 1990.

Gene H Golub and Victor Pereyra. The differentiation of pseudo-inverses and nonlinear least squares problems whose variables separate. SIAM Journal on numerical analysis, 10(2):413–432, 1973.

Christophe Geuzaine and Jean-François Remacle. Gmsh: A 3-d finite element mesh generator with built-in pre-and post-processing facilities. International journal for numerical methods in engineering, 79(11):1309–1331, 2009.

Anna Pisania, Gordon C Weir, John J O’neil, Abdulkadir Omer, Vaja Tchipashvili, Ji Lei, Clark K Colton, and Susan Bonner-Weir. Quantitative analysis of cell composition and purity of human pancreatic islet preparations. Laboratory investigation, 90(11):1661–1675, 2010.

Saba Parween, Elena Kostromina, Christoffer Nord, Maria Eriksson, Per Lindström, and Ulf Ahlgren. Intra-islet lesions and lobular variations in β-cell mass expansion in ob/ob mice revealed by 3d imaging of intact pancreas. Scientific reports, 6(1):1–11, 2016.

Johannes Reichold, Marco Stampanoni, Anna Lena Keller, Alfred Buck, Patrick Jenny, and Bruno Weber. Vascular graph model to simulate the cerebral blood flow in realistic vascular networks. Journal of Cerebral Blood Flow & Metabolism, 29(8):1429–1443, 2009.

Airlie Chapman. Advection on graphs. In Semi-Autonomous Networks, pages 3–16. Springer, 2015.

Radim Hošek and Jonáš Volek. Discrete advection–diffusion equations on graphs: Maximum principle and finite volumes. Applied Mathematics and Computation, 361:630–644, 2019.

Annie Rak. Advection on graphs. PhD thesis, 2017.

Diane T Finegood, Luisa Scaglia, and Susan Bonner-Weir. Dynamics of β-cell mass in the growing rat pancreas: estimation with a simple mathematical model. Diabetes, 44(3):249– 256, 1995.

Matthias König and M. Deepa Maheshvare. pancreas_minimal: Minimal model of pancreas glucose metabolism and insulin secretion. November 2021. doi: 10.5281/zenodo.5729745.

Marco Musy, Guillaume Jacquenot, Giovanni Dalmasso, neoglez, Ruben de Bruin, Ahinoam Pollack, Federico Claudi, Codacy Badger, icemtel, Bane Sullivan, Daniel Hrisca, Diego Volpatto, Nico Schlömer, Zhi-Qiang Zhou, and ilorevilo. marcomusy/vedo: 2021.0.2, March 2021.

Lee Mac. Converting a graph to a 2d diagram, 2020.

Ian S Kinstlinger, Sarah H Saxton, Gisele A Calderon, Karen Vasquez Ruiz, David R Yalacki, Palvasha R Deme, Jessica E Rosenkrantz, Jesse D Louis-Rosenberg, Fredrik Johansson, Kevin D Janson, et al. Generation of model tissues with dendritic vascular networks via sacrificial laser-sintered carbohydrate templates. Nature biomedical engineering, 4(9):916– 932, 2020.

Joshua R St Clair, David Ramirez, Samantha Passman, and Richard KP Benninger. Contrast-enhanced ultrasound measurement of pancreatic blood flow dynamics predicts type 1 diabetes progression in preclinical models. Nature communications, 9(1):1–12, 2018.

Juan D Rojas, Virginie Papadopoulou, Tomasz J Czernuszewicz, Rajalekha M Rajamahendiran, Anna Chytil, Yun-Chen Chiang, Diana C Chong, Victoria L Bautch, W Kimryn Rathmell, Stephen Aylward, et al. Ultrasound measurement of vascular density to evaluate response to anti-angiogenic therapy in renal cell carcinoma. IEEE Transactions on Biomedical Engineering, 66(3):873–880, 2018.

Oliver C Richards, Summer M Raines, and Alan D Attie. The role of blood vessels, endothelial cells, and vascular pericytes in insulin secretion and peripheral insulin action. Endocrine reviews, 31(3):343–363, 2010.

Elizabeth Haythorne, Maria Rohm, Martijn van de Bunt, Melissa F Brereton, Andrei I Tarasov, Thomas S Blacker, Gregor Sachse, Mariana Silva Dos Santos, Raul Terron Exposito, Simon Davis, et al. Diabetes causes marked inhibition of mitochondrial metabolism in pancreatic β-cells. Nature communications, 10(1):1–17, 2019.

Ronja M Malinowski, Seyed M Ghiasi, Thomas Mandrup-Poulsen, Sebastian Meier, Mathilde H Lerche, Jan H Ardenkjær-Larsen, and Pernille R Jensen. Pancreatic β-cells respond to fuel pressure with an early metabolic switch. Scientific reports, 10(1):1–11, 2020.

Ulrike Wittig, Renate Kania, Martin Golebiewski, Maja Rey, Lei Shi, Lenneke Jong, Enkhjargal Algaa, Andreas Weidemann, Heidrun Sauer-Danzwith, Saqib Mir, et al. Sabio-rk—database for biochemical reaction kinetics. Nucleic acids research, 40(D1):D790–D796, 2012.

Antje Chang, Lisa Jeske, Sandra Ulbrich, Julia Hofmann, Julia Koblitz, Ida Schomburg, Meina Neumann-Schaal, Dieter Jahn, and Dietmar Schomburg. Brenda, the elixir core data resource in 2021:new developments and updates. Nucleic Acids Research, 49(D1):D498–D508, 2021.

Yousef El-Gohary and George Gittes. Structure of islets and vascular relationship to the exocrine pancreas. Pancreapedia: The Exocrine Pancreas Knowledge Base, 2018.

M. Deepa Maheshvare, Soumyendu Raha, and Debnath Pal. A graph-based framework for multiscale modeling of physiological transport. bioRxiv, 2021. doi: 10.1101/2021.09.14.460337.

